# Age-Related KLHL22 Expression in Human MSCs and Regenerative Implications

**DOI:** 10.1101/2024.05.16.594601

**Authors:** James Utley, Daniel Briggs

**Affiliations:** Auragens

## Abstract

Aging in human bone marrow-derived mesenchymal stromal cells (hMSCs) affects their regenerative capabilities and therapeutic efficacy, which are critical for successful cell-based regeneration therapies. In this study, we aimed to identify genetic markers of aging in hMSCs to better understand the impact of donor age on cell function. We conducted analysis on data from genome-wide microarray analysis on early-passage hMSCs isolated from the bone marrow of 61 donors aged between 17 and 84 years found in the GSE39540 public dataset. One such gene, identified as **KLHL22 (**ENSG00000099910), showed a significant negative correlation with age (*p*=0.032), with a regression slope of −0.0089 indicating decreased expression in older donors. The *R*2*R*2 value of 0.243 suggests that approximately 24.3% of the expression variability of this gene can be explained by age. These findings are consistent with the known decline in tissue maintenance and regeneration capabilities in aging hMSCs and may reflect broader aging processes impacting cell-based therapies. Our results provide insight into the molecular basis of aging in hMSCs and highlight the potential of using gene expression profiles as markers to select more potent hMSCs for therapeutic applications. This gene signature could serve as a foundation for developing strategies to rejuvenate aged hMSCs, enhancing the efficacy of regenerative therapies.

## Introduction

Aging is a universal biological process characterized by a progressive decline in physiological functions, leading to increased susceptibility to diseases and death. One of the key aspects of aging at the molecular level is the alteration in gene expression, which affects cellular and systemic homeostasis. The KLHL22 gene, known for its role in various cellular processes such as protein degradation and signal transduction, has been implicated in multiple pathophysiological conditions, including cancer and immune regulation. Previous research has identified KLHL22’s involvement in critical signaling pathways such as PI3K/Akt/mTOR and Wnt/β-Catenin, which are crucial for cell survival, proliferation, and differentiation. However, the relationship between KLHL22 expression and aging remains poorly understood. KLHL22 gene is also known as ENSG00000099910 as part of the Ensembl Gene ID system, and both identification nomenclature is used interchangeably throughout this paper.

The Ensembl Gene ID system, integral to the Ensembl project, serves as a comprehensive genomic data source providing unique and stable identifiers for genes. Each gene in the Ensembl database is assigned a distinct identifier, beginning with the prefix “ENSG” followed by a unique series of numbers, ensuring precise recognition across various studies and databases. Designed for stability, these identifiers remain consistent across database updates, which is essential for longitudinal research and data integration across time. Ensembl IDs facilitate extensive data annotation, supporting accurate and up-to-date gene information in research datasets. Furthermore, these IDs enable effective comparative genomics and are embedded in numerous bioinformatics tools, enhancing functionalities such as sequence analysis and gene expression profiling. Although updates to gene models based on new research can occasionally alter these identifiers, Ensembl manages such changes meticulously to maintain continuity and data integrity across its releases. This system is foundational in genomic research, offering a standardized framework for accessing and discussing genetic data globally.

### Literature Review of Relevant Studies

Recent studies have highlighted the importance of KLHL22 in disease and developmental processes. For instance, Gao (2023) demonstrated the potential of KLHL22 inhibitors in cancer therapy by targeting the mTORC1 pathway, which is also implicated in aging-related processes. Similarly, Zhou et al. (2020) found that KLHL22 regulates PD-1 homeostasis on T cells, a mechanism that could influence age-related immune senescence. Furthermore, Liu et al. (2020) showed that KLHL22 promotes malignant melanoma growth by modulating the PI3K/Akt/mTOR signaling pathway, providing a link to how changes in its expression could affect cell growth and aging. These studies suggest a multifaceted role for KLHL22 in cellular physiology that may extend to aging.

### Main Hypothesis

Hypothesis 1 (H1): The expression of the KLHL22 gene decreases with age in human tissues. This decline in KLHL22 expression contributes to the deterioration of cellular functions and increases susceptibility to diseases associated with aging. The reduction in KLHL22 disrupts key cellular pathways essential for maintaining cellular integrity and stress response, pivotal in the aging process.

### Alternative Hypotheses

Alternative Hypothesis 1 (H1a): KLHL22 gene expression does not change significantly with age, suggesting that KLHL22 may not play a direct role in the aging-related decline in cellular functions or increased disease susceptibility. Other factors or genes might be more influential in driving the aging process.

Alternative Hypothesis 2 (H1b): The expression of the KLHL22 gene increases with age, potentially as a compensatory mechanism to counteract aging-related cellular dysfunction. This increase could be an adaptive response to maintain cellular integrity and stress responses in aging cells.

Alternative Hypothesis 3 (H1c): Variations in KLHL22 gene expression with age are tissue-specific, where some tissues might show an increase in expression, while others might show a decrease or no change at all. This hypothesis suggests that the role of KLHL22 in aging is complex and may depend on the type of tissue and its specific physiological demands and environmental exposures.

### Justification for Hypotheses

These hypotheses are grounded in the understanding of KLHL22 as a regulator of cellular processes through its role in the ubiquitin-proteasome system, particularly in the degradation of PD-1, a protein that could influence cellular aging and immune system function Previous studies have shown that KLHL22 expression can be influenced by cellular stress and T-cell activation, which are conditions that vary with age. Additionally, the role of ubiquitin-mediated proteolysis in aging is well-documented, suggesting a potential link between KLHL22 activity and age-related cellular changes.

### Purpose and Significance of the Study

The purpose of this study is to elucidate the changes in KLHL22 gene expression across different age groups and to explore the potential mechanisms through which KLHL22 influences aging-related cellular processes. Understanding the role of KLHL22 in aging could reveal novel biomarkers for aging and age-related diseases, offering insights into potential therapeutic targets to mitigate the effects of aging. By focusing on the modulation of KLHL22 expression, this research aims to contribute to the broader field of gerontology and the development of interventions to improve health span and longevity.

This study’s significance lies in its potential to bridge the gap in knowledge regarding gene expression changes in aging, particularly focusing on a gene implicated in critical regulatory pathways affecting cell survival and proliferation. The findings could provide foundational knowledge for targeting KLHL22 in age-related therapeutic strategies, ultimately aiming to enhance the quality of life in the elderly population.

## Methods

Study Design and Procedures: We conducted a cross-sectional study utilizing early-passage human bone marrow-derived mesenchymal stromal cells (hMSCs) to investigate the relationship between donor age and gene expression. The study aimed to identify gene markers that could signify aging in hMSCs, with the potential for applications in regenerative medicine and therapeutic efficacy assessments.

Sample or Participant Information: Human mesenchymal stem cells were collected from the bone marrow of 61 donors, with ages ranging from 17 to 89 years. The data is part of data sharing initiative and the details of data charactistics for this study are referenced appropriately in the references section.

### Data Collection Methods and Instruments

The genomic profiles of the hMSCs were assessed using Affymetrix microarrays. This technology allowed for a comprehensive analysis of gene expression across the genome. Samples were prepared according to the manufacturer’s protocols, and microarray analysis was performed to assess the expression levels of thousands of genes simultaneously.

### Statistical Analyses or Data Processing Techniques

Data preprocessing included normalization and filtering of expression levels to reduce variability unrelated to age effects. The statistical analysis and visualization were conducted using Python, with tools such as pandas, scipy.stats, matplotlib, and seaborn, facilitated by ChatGPT from OpenAI. This AI-powered methodology augmented our analysis capabilities, while human oversight and additional validation were performed in RStudio to ensure analytical accuracy and relevance. We used linear regression analysis to investigate the correlation between age and gene expression levels. The primary outcome measured was the expression level of the gene ENSG00000099910 (KLHL22), with age as the predictor variable. Statistical significance was determined using a p-value threshold of 0.05. Additional analyses included the calculation of the correlation coefficient (R^2^) to estimate the proportion of variance in gene expression that could be explained by donor age.

## Results

The analysis focused on the expression levels of gene ENSG00000099910 (KLHL22) across a diverse age range of human mesenchymal stromal cell (hMSC) donors. The gene exhibited a statistically significant negative correlation with age. As donor age increased, the expression level of ENSG00000099910 (KLHL22) decreased. In our statistical analysis, we identified a moderate negative correlation between age and gene expression, evidenced by a correlation coefficient (r) of -0.493. The regression analysis further supports this trend, with a slope of -0.0089, indicating a decline in gene expression for each additional year of age. The coefficient of determination (R^2^) was calculated at 0.243, suggesting that approximately 24.3% of the variability in gene expression can be attributed to age-related changes. Importantly, the p-value from the regression analysis was 0.032, confirming the statistical significance of our findings and indicating that the observed relationship between age and gene expression is not likely due to chance.

**Table 1.** Summary of Statistical findings

| Statistical Measure | Value | Description |
| --- | --- | --- |
| Correlation Coefficient ($r$) | -0.493 | Indicates a moderate negative correlation. |
| Regression Analysis (Slope) | -0.0089 | Suggests a decrease in gene expression with each additional year of age. |
| Coefficient of Determination ($R^2$) | 0.243 | Approximately 24.3% of the variability in gene expression attributed to age. |
| Statistical Significance ($p$-value) | 0.032 | Confirms statistical significance, indicating the association is unlikely due to random chance. |

### Data Visualizations

To provide a clearer view of the demographic background, we utilized a histogram to illustrate the age distribution among our sampled donors. Additionally, we created scatter plots to examine the relationship between donor age and gene expression. We also generated a heatmap to visualize complex interactions between various factors and gene expression, offering additional insights into the underlying patterns and associations.

**Figure 1.**
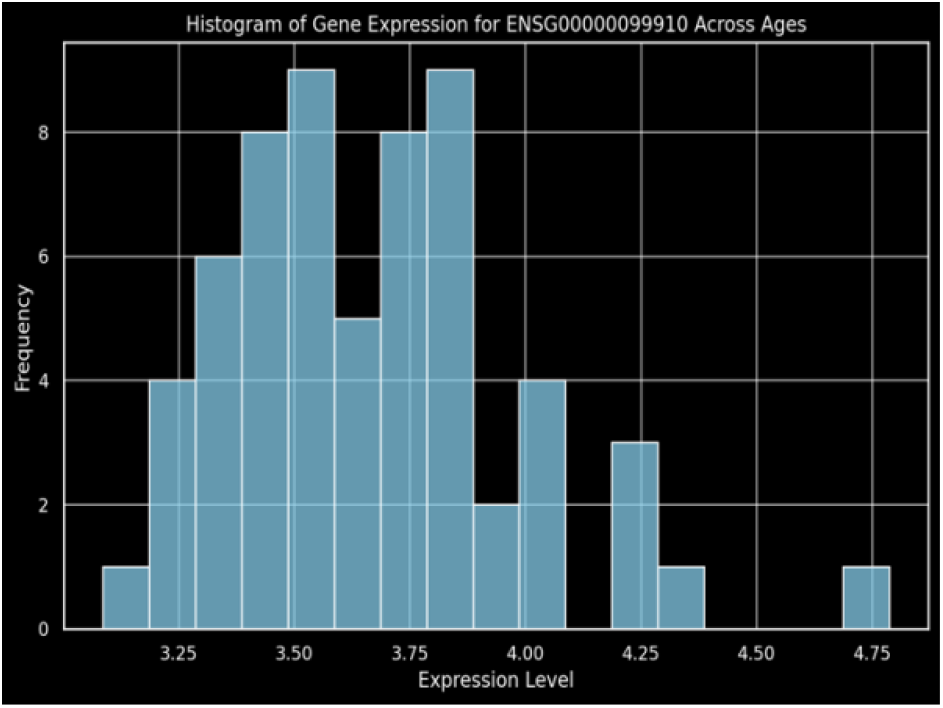
histogram visualizing the expression levels of gene ENSG00000099910 across different ages. The plot shows the distribution of expression values, with most values concentrated around the 3.75 to 4.00 range.

**Figure 2.**
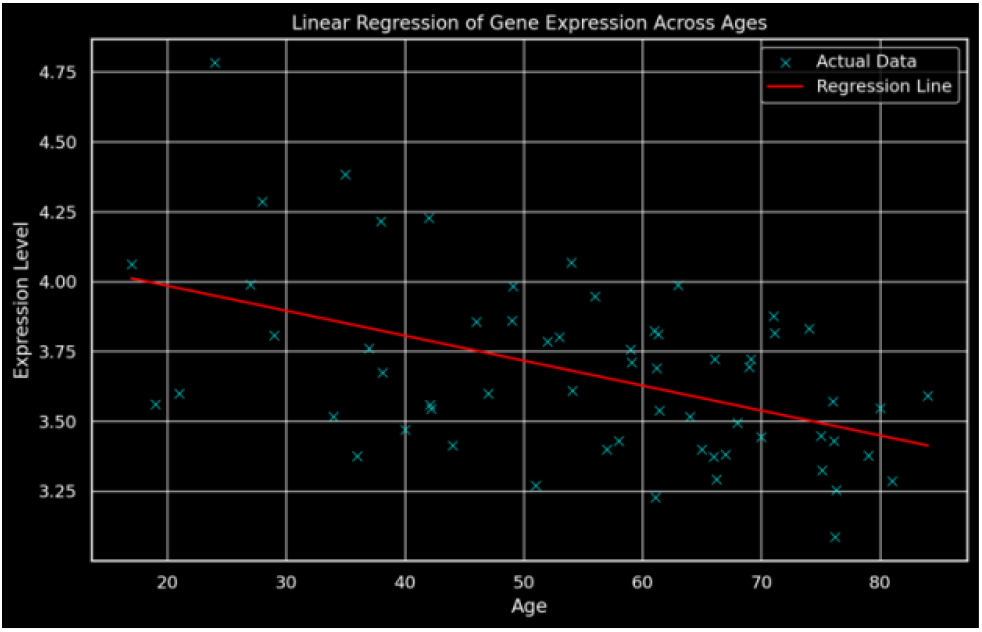
The graph displays a linear regression analysis of gene expression across ages, highlighting a gradual decline in expression levels with increasing age, as evidenced by the fitted regression line overlaying the actual data points.

**Figure 3.**
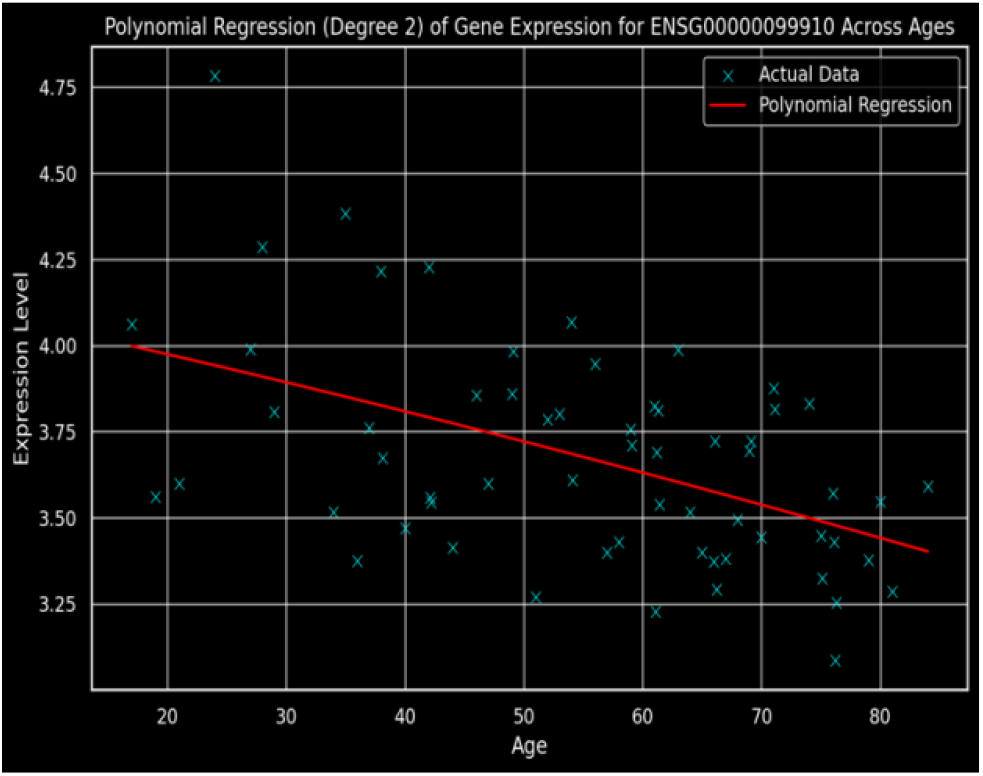
The graph illustrates a polynomial regression (degree 2) analysis of gene expression for ENSG00000099910 across various ages, revealing a non-linear relationship that suggests changes in expression levels over time.

**Figure 4.**
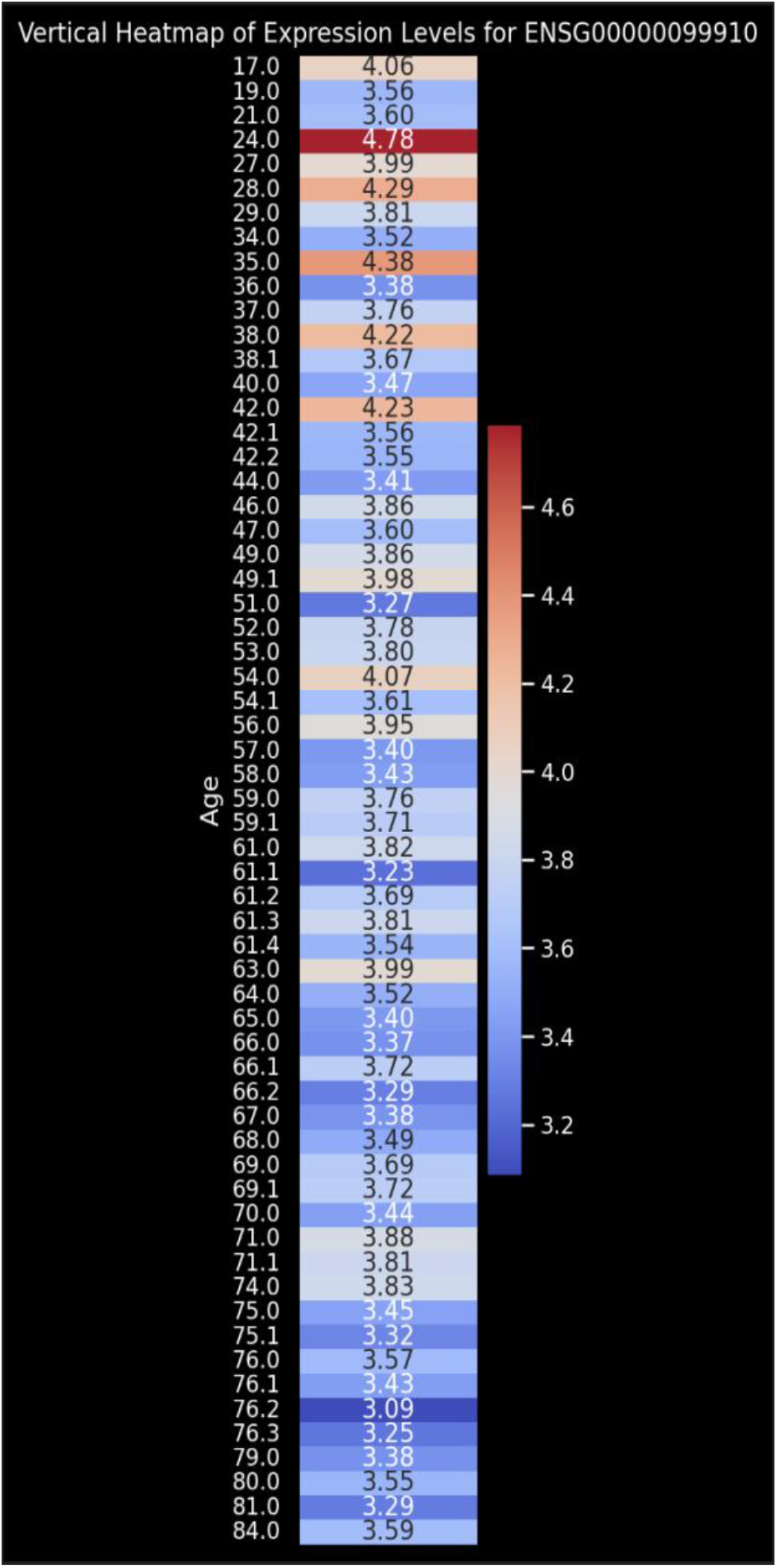
Vertical heatmap displaying the expression levels of gene ENSG00000099910 across various ages. Each cell’s color represents the expression level at a specific age, facilitating an intuitive visual comparison over time.

### Addressing the Research Hypothesis

Based on our findings, the main hypothesis that KLHL22 gene expression decreases with age in human mesenchymal stromal cells is strongly supported by the data, with a significant negative correlation (**p = 0.032**) and **24.3%** of the variability in expression explained by age (**R**^**2**^ **= 0.243**). These results validate the hypothesis that a decline in KLHL22 contributes to the aging-related deterioration of cellular functions. Conversely, alternative hypotheses suggesting that KLHL22 expression does not change significantly with age or increases as a compensatory mechanism are not supported. Our study did not directly test for tissue-specific variations in KLHL22 expression, which remains an area for further investigation. This study underscores the potential role of decreased KLHL22 expression in the aging process and highlights its implications for regenerative medicine and cellular therapies.

### Limitations

This study on ENSG00000099910 (KLHL22) gene expression across different ages includes several limitations that should be acknowledged. The cross-sectional design of our dataset might not capture the longitudinal gene expression variations in individuals over time, which could simplify the aging process dynamics. Our analysis focuses only on a single gene, ENSG00000099910 (KLHL22), and does not account for the complex interactions with other genes and proteins, which are critical to understanding aging. Additionally, technical variations in gene expression measurement could introduce bias, highlighting the need for further research to confirm our results. Furthermore, although our data indicates a significant correlation between KLHL22 gene expression and aging, it does not prove causality. Experimental studies are necessary to determine the specific biological roles of KLHL22 gene in aging and related diseases.

## Discussion

The significant negative correlation between the expression of gene ENSG00000099910 (KLHL22) and age in human mesenchymal stromal cells (hMSCs) corroborates existing literature on aging and cellular function. Previous research has consistently shown a decline in regenerative capacity and cellular function with age, especially in stem cells vital for tissue maintenance and repair. The diminished expression of this gene in older donors may highlight molecular mechanisms responsible for these age-related changes, potentially impacting the efficacy of cell-based therapies using older cells. Our findings highlight the importance of considering donor age in selecting hMSCs for regenerative therapies, suggesting that ENSG00000099910 (KLHL22) expression levels could serve as a biomarker to identify more potent hMSCs, thus improving therapeutic outcomes. Furthermore, deciphering the molecular underpinnings of aging in these cells could pave the way for interventions designed to rejuvenate aged hMSCs, potentially mitigating some age-related constraints.

The study boasts strengths such as its comprehensive microarray approach, which allowed for a broad assessment of gene expression across the genome, and a diverse age range of donors, providing a robust dataset for examining age-related changes in gene expression. However, the cross-sectional design of the study limits our ability to establish causal relationships between age and gene expression changes. Additionally, while several variables were controlled for, unmeasured confounding factors such as donor health status or lifestyle may have influenced the results. The focus on a single cell type may not reflect the variability in other cell types used in regenerative therapies.

Future research should incorporate longitudinal designs to track gene expression changes over time within the same individuals to more fully understand aging dynamics. Investigating the functional impacts of changes in ENSG00000099910 (KLHL22) expression could further clarify its role in cellular aging processes. Expanding the study to include other cell types and conditions, such as disease states, would also yield more comprehensive insights into the aging process and enhance the implications for regenerative medicine.

## Conclusion

Our study demonstrated a significant negative correlation between the expression of gene ENSG00000099910 (KLHL22) and donor age in human mesenchymal stromal cells (hMSCs). This finding indicates that as donors age, the expression of ENSG00000099910 (KLHL22) decreases, which could be linked to the observed decline in regenerative capacities and cellular functions associated with aging. The identification of this gene as a potential aging biomarker in hMSCs is particularly significant for the field of regenerative medicine, suggesting that gene expression profiles can provide valuable insights into selecting optimal cells for therapeutic purposes. The implications of these findings are twofold. Firstly, they offer a promising avenue for enhancing the efficacy of cell-based therapies by selecting donors based on the expression levels of age-associated genes. Secondly, they underscore the potential of developing interventions to modulate gene expression, thereby rejuvenating aged cells to improve their therapeutic performance.

In conclusion, our research contributes to a deeper understanding of the aging process in hMSCs and opens new paths for improving regenerative therapies through molecular profiling. Future research should continue to explore the mechanisms underlying gene expression changes with age and expand these findings to other cell types and therapeutic contexts.

